# Differential Nucleotide Inhibition Profile of Mouse and Human UCP1 Expressed in Liver Mitochondria Is Associated with an F88S Mutation

**DOI:** 10.64898/2026.08.19.745785

**Authors:** Irina G. Shabalina, Luise Jacobsen, Glauber Rudá Feitoza Braz, Zhi Wei Zeng, Qimuge Naren, Björn Eriksson, Uzair Ali, Jiaping Li, Annika Ericsson, Barbara Cannon, Himanshu Khandelia, Jan Nedergaard

**Author notes:** Corresponding author: Jan Nedergaard, The Department of Molecular Biosciences, The Wenner-Gren Institute, Stockholm University, SE-106 91 Stockholm, Sweden.

## Abstract

Uncoupling protein 1 (UCP1) mediates thermogenesis in brown adipose tissue. Whether human-UCP1 shares the bioenergetic properties established for rodent UCP1 (innate uncoupling, GDP sensitivity, fatty acid (re)activation) is not known. Therefore, we expressed human and mouse UCP1 in mouse liver, using adeno-associated viral vectors, and characterized their properties in isolated liver mitochondria. Both UCP1s induced marked innate uncoupling, characterized by increased substrate-supported respiration and decreased membrane potential, in the absence of exogenous fatty acids. Mouse-UCP1 in liver retained the classical regulatory properties of native brown-fat UCP1, including potent inhibition by GDP and reactivation by oleate. In contrast, human-UCP1 was only weakly inhibited by GDP but was strongly responsive to fatty acids. However, ATP potently inhibited human-UCP1, with an apparent IC₅₀ of ≈0.4 mM compared with ≈1.4 mM for GDP, and ATP markedly decreased the sensitivity of human-UCP1 to oleate (re)activation. Despite substantial UCP1-mediated uncoupling, oxidative phosphorylation capacity and mitochondrial OXPHOS protein levels were preserved. Molecular dynamics simulations suggested a structural basis for the species difference. GDP formed persistent interactions with F88 in mouse-UCP1, an interaction absent at the corresponding S88 residue in human-UCP1. In-silico substitution of F88 by serine reduced GDP interaction at this site. Thus, human and mouse UCP1 share innate thermogenic activity but differ fundamentally in nucleotide regulation. The F88/S88 difference may contribute to the preferential GDP sensitivity of mouse-UCP1, whereas ATP provides effective nucleotide control of human-UCP1.

**Highlights:**

- Human and mouse UCP1 expressed in liver mitochondria are innately uncoupling active
- Human and mouse UCP1 in liver confers high fatty acid uncoupling sensitivity
- Mouse UCP1 in liver retains strong GDP inhibition
- Human UCP1 in liver is poorly inhibited by GDP but potently inhibited by ATP
- F88 in mouse UCP1 stabilizes GDP binding in molecular dynamic simulations
- The F88S substitution may underlie the species-specific nucleotide regulation

## INTRODUCTION

Since it became generally accepted that not only newborn humans but also adult humans possess physiologically active brown adipose tissue (Nedergaard et al., 2007, Cypess et al., 2009, van Marken Lichtenbelt et al., 2009, Virtanen et al., 2009, Saito et al., 2009), the interest specifically for human brown adipose tissue has greatly expanded. This expansion includes evidently understanding the function of human UCP1, as compared to earlier studies of UCP1 that mainly have been performed in rodent UCP1. Indeed, the structure of human UCP1 has now been known for several years (Kang and Chen, 2023, Jones et al., 2023), but many basic functional features of this unique protein remain mechanistically unsolved.

One prominent such feature is whether the high proton conductivity (that is the background for the thermogenic function of UCP1 and thus for the thermogenic function of brown adipose tissue (Nicholls and Locke, 1984, Cannon and Nedergaard, 2004)) is an innate property of UCP1 – or whether it is induced by in-situ components. Particularly, in reconstituted systems (e.g. liposomes or black lipid membranes), UCP1 is not associated with a high innate proton conductivity (Winkler and Klingenberg, 1994, Garlid et al., 1996, Urbánková et al., 2003, Cavalieri et al., 2022). The high conductivity observed in brown-fat mitochondria could therefore be a function of e.g. the particular mitochondrial membranes in brown adipose tissue.

In this context, it is important that it has been suggested that human UCP1 is divergent from e.g. mouse UCP1 in that this high innate proton conductivity may not be seen in human UCP1. This could be understandable, given that out of the 307 amino acids that constitute UCP1, as many as 64 are different between mouse and human UCP1, allowing for species-specific properties. There have also been observations implying that the nucleotide regulation is divergent between human and mouse UCP1 (Musiol et al., 2024).

To approach several of these outstanding questions, we have developed a system for ectopic expression of human- and mouse-UCP1 in mouse liver mitochondria through adeno-associated virus (AAV) gene delivery. In the present study, we investigate the bioenergetic characteristics of ectopically expressed mouse-UCP1 and human-UCP1. We assess the bioenergetic impact of these UCP1s through measuring oxygen consumption, membrane potential, and responsiveness to fatty acids and to nucleotides. We uncover distinct binding behaviour of GDP and ATP between human-UCP1 and mouse-UCP1, and we explain these differences through molecular dynamics simulations.

## MATERIALS AND METHODS

### Animals and AAV injection

Male C57BL/6J mice (10–12 weeks old) were obtained from Charles River Laboratories and housed in groups of 4–5 per cage, with environmental enrichment, at 21 °C (room temperature) under a 12-hour light/dark cycle. Mice were provided with standard chow (Altromin 1324; Brogaarden, Sweden) and water ad libitum. After an acclimatization period of at least two weeks, animals received intravenous injections via the tail vein with adeno-associated viral vectors (AAV, serotype 8), diluted in phosphate-buffered saline. The following vectors were used as indicated: a control empty vector (AAV-Empty); a mouse UCP1 (mUCP1) vector driven by the ubiquitous CAG promoter (AAV8-CAG-mUCP1); a human UCP1 (hUCP1) vector under the same CAG promoter (AAV8-CAG-hUCP1); and a human UCP1 vector regulated by a liver-specific composite element, consisting of the Serpina 1 enhancer CRM8 (Chuah et al., 2014) and the TTR (Samadani et al., 1996) (AAV8-CRM8/TTR-hUCP1). All vectors were administered at a dose of 2 × 10¹³ genome copies (gc) per kg body weight, corresponding to an injection volume of 2 µl per g body weight. All vectors were produced by VectorBuilder (China). The UCP1 mRNA sequence was optimized for expression.

Four to six weeks after the AAV injections, the mice were euthanized by CO₂ inhalation. Livers were rapidly excised and weighed. One small piece of liver tissue was snap-frozen in liquid nitrogen for later mRNA analysis. The remaining liver tissue was immediately immersed in ice-cold mitochondrial isolation buffer for isolation of mitochondria for respirometry and western blotting.

All animal procedures were approved by the Local Stockholm Animal Experiment Ethics Committee.

### RNA preparation, cDNA synthesis, and quantitative RT-qPCR

Total RNA was isolated using TRIzol Reagent™ (Invitrogen) according to the manufacturer’s protocol. Chloroform (200 µl) was added to each sample, followed by vigorous shaking and centrifugation at 14 000 g for 30 min at 4 °C. The aqueous phase was collected and mixed with an equal volume of isopropanol. After incubation on ice for 10 min, the samples were centrifuged at 12 000 rpm for 60 min at 4 °C. The RNA pellet was washed with 1 ml ice-cold 75% ethanol, centrifuged at 14 000 g for 20 min at 4 °C, air-dried, and resuspended in 1 mM EDTA. RNA concentrations were measured using a NanoDrop 2000 spectrophotometer (Thermo Scientific).

cDNA was synthesized using the High-Capacity cDNA Reverse Transcription Kit (Applied Biosystems). Each reaction contained 10 µl (500 ng) of RNA, random hexamer primers, dNTPs, MultiScribe reverse transcriptase, and an RNase inhibitor. Quantitative PCR (qPCR) was performed using 2 µl of cDNA and Maxima Probe/ROX qPCR Master Mix (Thermo Scientific), along with gene-specific primers and DEPC-treated water. Reactions were run in triplicate on a CFX Connect™ Real-Time PCR System (Bio-Rad). Gene expression was quantified using the ΔΔCt method and normalized to eukaryotic 18S rRNA.

TaqMan™ Gene Expression Assays were from Applied Biosystems. The primer and probe sequences used were: for mUCP1, forward primer GCAATCTCACCTGCATGGTATTAAG, reverse primer GTACTGAGACTTTCTGTGGTAGCAA, and probe CCGAGATATACAGGAACATAC; for hUCP1, forward primer AAGCCTGGGTAGCAAGATTC, reverse primer AGATGGGACTGTGCTTGGAG, and probe TTGCTGGACTGACAACTGG.

### Mitochondrial isolation

Liver mitochondria were isolated principally as earlier described (Gómez Rodríguez et al., 2022). Liver tissue was finely minced with scissors and homogenized in ice-cold buffer composed of 70 mM sucrose, 210 mM mannitol, 20 mM TES, 1 mM EDTA, and 0.2 % fatty-acid-free bovine serum albumin (BSA) (in some experiments, the BSA concentration was increased to 0.6 %, as indicated). The homogenate was centrifuged at 8800 g for 10 min at 4 °C, and the supernatant, including the upper fat layer, was discarded. To remove nuclear debris, the pellet was resuspended in fresh isolation buffer and centrifuged at 800 g for 10 min at 4 °C. To collect the mitochondrial pellet, the resulting supernatant was centrifuged again at 8800 g for 10 min at 4 °C. This pellet was washed by resuspension in the same buffer and then centrifuged again at 8800 g for 10 min. The final mitochondrial pellet was resuspended in the same medium using a small glass homogenizer. Protein concentration was determined using the fluorescamine method with BSA as the standard (Udenfriend et al., 1972).

### Western blotting

Thawed mitochondrial samples were re-quantified using the Lowry method and mixed with an equal volume of reducing buffer (0.5 M Tris-HCl, pH 6.8, 10 % SDS, 2.5 % glycerol, 100 mM DTT, and 0.5 % bromophenol blue). Samples (10–20 µg protein) were separated on 12 % SDS-PAGE gels. Proteins were transferred to PVDF membranes (GE Healthcare) in transfer buffer (48 mM Tris-HCl, 39 mM glycine, 0.037 % SDS, 15 % methanol) using a semi-dry blotting system (Bio-Rad) at 1.2 mA/cm² for 90 min. Membranes were blocked for 1 h in 5 % milk in TBST and incubated overnight at 4 °C with primary antibodies.

Mouse-UCP1 was detected using a rabbit antiserum raised against the C-terminal decapeptide of mouse-UCP1, diluted 1:10 000 (Shabalina et al., 2013). Human UCP1 was detected using a rabbit polyclonal antibody from Abcam (ab10983), directed against a synthetic peptide corresponding to human-UCP1 residues 145–159 conjugated to keyhole limpet hemocyanin (1:5000). For mitochondrial complex detection, the Total OXPHOS Rodent Antibody Cocktail (MS601, Mitosciences) was used at 1:1000, and HADHA served as the loading control. Signal detection was performed using horseradish peroxidase-conjugated secondary antibodies and ECL (GE Healthcare), visualized by ChemiDoc (Bio-Rad), and quantified with ImageJ software.

To produce the hUCP1 used here as a positive control for protein size, the human-UCP1 coding sequence had been cloned into a pcDNA3.1^+^ vector, amplified, and transfected into HEK293 cells (Jastroch et al., 2012). Protein lysate (0.5 µg) from these cells was loaded on each gel as a reference. Brown adipose tissue protein extracts from C57BL/6J mice room temperature-housed were used as a positive control for mouse-UCP1.

### Mitochondrial oxygen consumption

Oxygen consumption was measured at 37 °C using the Oxygraph-2k system (Oroboros Instruments) as described previously (Shabalina et al., 2025). Mitochondria (0.5 mg protein) were suspended in incubation buffer containing 100 mM sucrose, 20 mM K-TES (pH 7.2), 50 mM KCl, 4 mM KH₂PO₄, 2 mM MgCl₂, 1 mM EDTA, and 0.1 % fatty-acid-free BSA. Basal respiration was measured after adding 6 mM malate, 10 mM glutamate, and 10 mM pyruvate. Octanoyl-L-carnitine (0.5 mM) was added in some assays, as indicated.

ADP (1 mM) was added to stimulate oxidative phosphorylation. Oligomycin (3 µg/ml) or carboxyatractyloside (CATR) (3 µM) was used to inhibit ATP synthase or ATP/ADP- antiporter, respectively. Maximal respiration was measured after titration with FCCP (0.8 µM). Oleate was added in stepwise concentrations (10–20 µM) in the presence of CATR. ATP dose-response was studied in the presence of both oligomycin and CATR. Rates were calculated as means over 1 min intervals.

### Data analysis for inhibitor and activator response curves

IC_50_ values for nucleotide-induced inhibition and EC_50_ values for fatty acid activation of UCP1- dependent respiration were calculated by nonlinear regression. Concentration-response curve data were analyzed with the general fit option of the KaleidaGraph version 5.01 by Synergy Software (Reading PA, USA) for adherence to the Hill equation. Apparent EC_50_ and IC_50_ values and the degree of cooperativity of the system (the Hill coefficient) were estimated using the equation V(x) = Vbasal + (Vmax-Vbasal) • (x^Hill/(EC50^Hill + x^Hill)), or similarly for inhibition V(x) = Vmax - Vbasal • (x^Hill/(IC50^Hill + x^Hill)).

Oleate dose-response curves were evaluated under both nominal and free fatty acid concentrations. To estimate free [oleate], the binding of oleate to bovine serum albumin (BSA) was modeled using a known partition coefficient (mol:mol ratio of binding sites per BSA molecule) as previously described in (Shabalina et al., 2004). The total added oleate and BSA concentration were used to calculate free fatty acid levels via equilibrium binding equations.

### Mitochondrial membrane potential

For mitochondria from mUCP1-expressing animals, TMRM (0.5 µM) fluorescence was measured in parallel with respiration at 37 °C using the O2k-Fluo LED2 module (Shabalina et al., 2025). Calibration was performed by stepwise titration of KCl (0.1–100 mM) in the presence of valinomycin (3 µM), and complete depolarization was induced by alamethicin (10 µM). Δψ was calculated using the Nernst equation: Δψ = 61 mV × log([K⁺]in/[K⁺]out).

For mitochondria expressing hUCP1 two methods assessing membrane potential were used: TMRM fluorescence as above and safranin O absorbance at 511–533 nm using an Olis Aminco DW-2 spectrophotometer (Nedergaard, 1983). Calibration and calculation followed the same K⁺ titration and valinomycin protocol.

### Chemicals

The following chemicals were used: fatty-acid-free bovine serum albumin (BSA), Fraction V (Cat#10775835001, Roche Diagnostics GmbH); malate (sodium salt) (Cat#M9138, Sigma-Aldrich); pyruvate (sodium salt) (Cat#P2256, Sigma-Aldrich); glutamate (monosodium salt) (Cat#G5889, Sigma-Aldrich), octanoyl-L-carnitine (Cat#50892, Sigma-Aldrich) and the adenine nucleotide translocase inhibitor carboxyatractyloside (CATR) (Cat#216200, Calbiochem), all dissolved in water. ADP (sodium salt) (Cat#A2754, Sigma-Aldrich), ATP (disodium salt) (Cat# A2383, Sigma-Aldrich) and GDP (sodium salt) (Cat#G7127, Sigma-Aldrich) were dissolved in 20 mM TES (pH 7.2). Oligomycin (Cat#O4876, Sigma-Aldrich), FCCP (carbonyl cyanide 4-(trifluoromethoxy)phenylhydrazone) (Cat#C2920, Sigma-Aldrich), valinomycin (Cat#V0627, Sigma-Aldrich) and alamethicin (Cat#A4665, Sigma-Aldrich) were dissolved in 95% ethanol. Oleate (sodium salt) (Cat#O7501, Sigma-Aldrich) was dissolved in 50 % ethanol. TMRM (tetramethylrhodamine methyl ester) (Cat#T5428, Sigma-Aldrich) was dissolved in DMSO. Final ethanol or DMSO concentrations were verified to have no effect on mitochondrial function. The remaining chemicals used for mitochondrial isolation and respiration media were of GC purity >99% and were obtained from Sigma-Aldrich. Fluorescamine (Cat# F9015, Sigma-Aldrich) was dissolved in acetone according to protein determination protocol (Udenfriend et al., 1972).

### Statistics

Data were analyzed using Prism 4 (GraphPad) or KaleidaGraph 5.0 (Synergy Software). Values are presented as means ± SEM. Comparisons between groups were made using Student’s *t*-test or one-way ANOVA. Statistical significance was accepted at p < 0.05.

### Computational UCP1 models

The computational mUCP1 model was obtained from AlphaFold (Jumper et al., 2021, Varadi et al., 2022), https://alphafold.ebi.ac.uk/entry/P12242 visited: November 6 2024] and the ATP-bound hUCP1 structure was taken from the Protein Data Bank (pdb 8hbw). The AlphaFold confidence (pLDDT) score was 74.86 for mUCP1, indicating high reliability. The root mean square deviation (RMSD) between the backbone of the mUCP1 AlphaFold model and the Cryo-EM structure of apo hUCP1 (pdb 8hbv) (Kang and Chen, 2023) was low (4.06 Å), further supporting the AlphaFold model’s accuracy. UCP1 was embedded in a membrane with a composition resembling an inner mitochondrial membrane (Daum and Vance, 1997, Horvath and Daum, 2013), and three cardiolipin molecules were pre-positioned at their established binding sites on UCP1 (Jacobsen et al., 2023, Kang and Chen, 2023, Jones et al., 2023).

### Molecular dynamics simulations

All simulations were performed using GROMACS 2021.1 (Pronk et al., 2013, Abraham et al., 2015) or newer, with the CHARMM36m force field (Brooks et al., 2009, Huang et al., 2017) . Molecular graphics were generated using VMD (Humphrey et al., 1996). ProLIF (Bouysset and Fiorucci, 2021) was used to compute the fraction of simulation time with nucleotide-protein interaction for each residue, and PyLipID (Song et al., 2022) was used to determine the final bound conformations.

The simulation systems were prepared using the membrane builder function in CHARMM-GUI (Jo et al., 2008, Lee et al., 2016, Park et al., 2021). However, nucleotide substitutions and ion neutralization were performed manually. Each simulation system contained ∼26,000 water molecules and 150 mM NaCl.

Energy minimization was carried out for 5,000 steps using the steepest descent algorithm, followed by a five-step equilibration process in which restraints on the system were gradually lifted. For each of the three systems (ATP-hUCP1, GDP-hUCP1, and GDP-mUCP1), three 1-1.4 µs production simulation runs were performed. The last 300 ns of each simulation were used for analysis.

The simulation time step was 2 fs and the pressure was maintained at 1 atmosphere using the semi-isotropic C-rescale barostat (Bernetti and Bussi, 2020) with a time constant of 5 ps and a compressibility of 0.00045/bar. The temperature was kept at 310 K using the V-rescale thermostat (Bussi et al., 2007) with a time constant of 1ps. The Linear Constraint Solver algorithm (Hess et al., 1997) was used to constrain all bonds involving hydrogen atoms. A 1.2 nm cut-off was applied for short-range interactions, and Particle Mesh Ewald (Darden et al., 1993, Essmann et al., 1995) was applied for long-range electrostatic interactions. Sequence alignment between human UCP1 (hUCP1, UniProtID: P25874) and mouse UCP1 (mUCP1, UniprotID: P12242) was performed using the UniProt alignment tool (Consortium, 2023).

## RESULTS

We have here first examined the properties of liver mitochondria isolated from mice with mouse-UCP1 expressed in the liver, and then liver mitochondria from human-UCP1-expressing mice. We have then through molecular dynamics identified structural changes that may explain the observed distinct regulatory properties of human-UCP1 versus mouse-UCP1.

### Mouse UCP1 is innately thermogenic even in an ectopic membrane environment

Innately protonophorically active UCP1 – and the inhibition of this innate UCP1 activity by GDP – are generally considered fundamental properties of brown-fat mitochondria, distinguishing brown-fat mitochondria from mitochondria from any other tissue (Cannon et al., 1973, Nicholls, 1976, Matthias et al., 1999, Monemdjou et al., 1999, Jimenez-Jimenez et al., 2006, Shabalina et al., 2010, Yu et al., 2023). However, this innate activity is, as mentioned, not observed in reconstituted systems. This raises the question of whether UCP1’s innate uncoupling activity depends on the specific environment of brown adipose tissue mitochondria.

To investigate whether the functional characteristics of UCP1 would be equally manifest in a liver mitochondrial environment, we injected mice with AAV8-CAG-mUCP1. About one month after the injection we compared the bioenergetics of these liver mitochondria to that of liver mitochondria from mice injected with AAV-Empty (referred to as controls).

Quantitative PCR analysis demonstrated high levels of mUCP1 mRNA in liver tissue (Figure 1a), and western blotting of isolated liver mitochondria revealed a specific immunoreactive band for mUCP1 at ∼33 kDa, that was absent in controls, as well as in brown-fat mitochondria from UCP1 knockout mice (Figure 1b).

**Figure 1.**
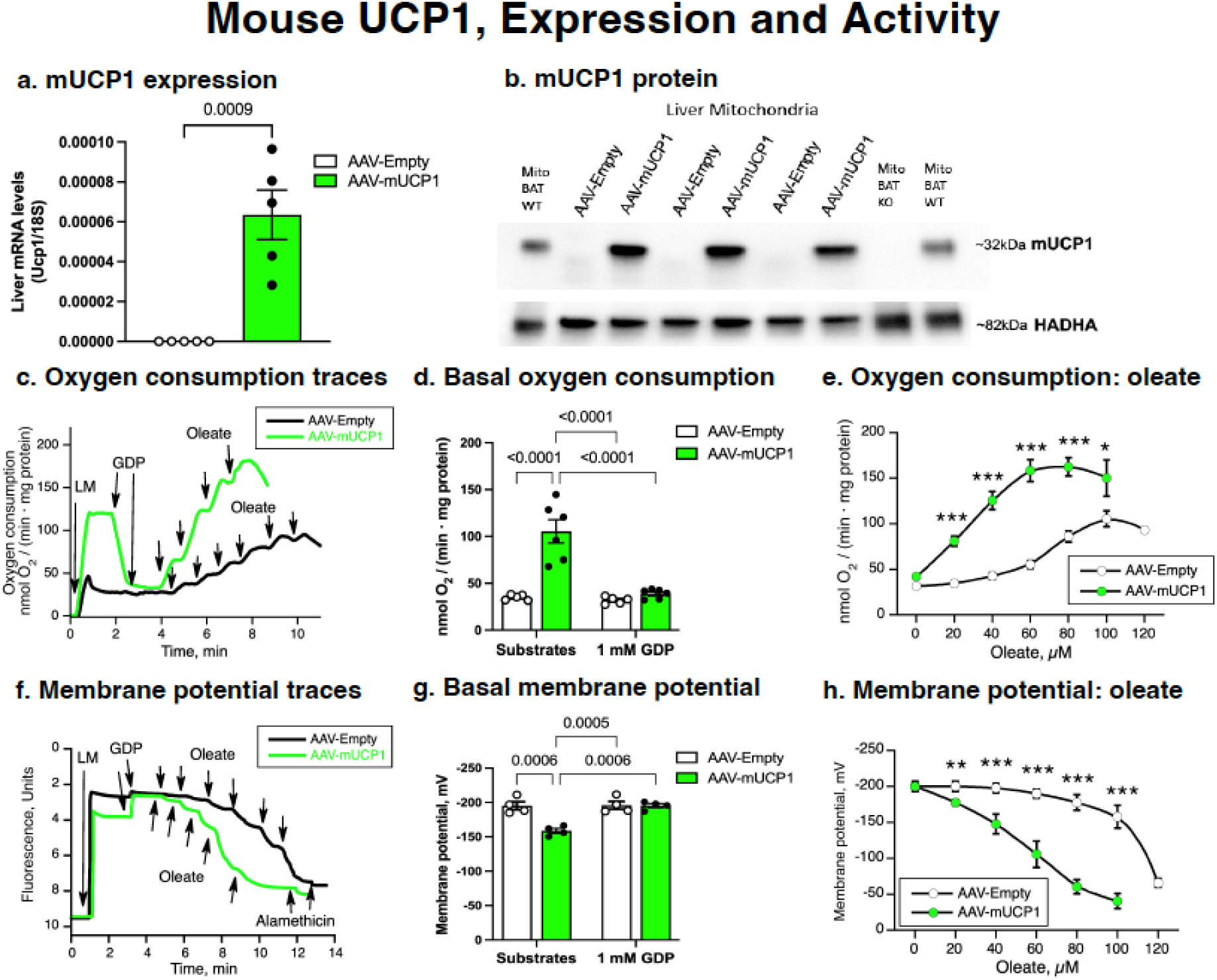
Mouse-UCP1 expression and activity in liver mitochondria. (a) Relative mRNA levels of mUCP1 in liver tissue of AAV8-CAG-mUCP1 or AAV-Empty-injected mice, normalized to 18S rRNA. (b) Western blot analysis of mUCP1 protein expression in isolated liver mitochondria; brown adipose tissue (BAT wild-type and UCP1 knockout (KO)) included as a positive control. HADHA was used as a loading control. (c) Representative traces of oxygen consumption in isolated liver mitochondria (0.5 mg protein) expressing mUCP1 or control, measured at 37 °C in the presence of substrates (6 mM malate, 10 mM glutamate, and 10 mM pyruvate), followed by the addition of 1 mM GDP and stepwise additions of 20 µM sodium oleate. (d) Basal oxygen consumption rate (innate UCP1 activity) measured in the presence of substrates and after addition of 1 mM GDP compiled from (c). (e) Oleate-stimulated oxygen consumption in the presence of 1 mM GDP. (f) Representative traces of mitochondrial membrane potential changes recorded by TMRM fluorescence following sequential addition of 0.5 mg liver mitochondria in the presence of substrates as in (c), 1 mM GDP, and titration with sodium oleate (each addition 20 µM). (g) Steady-state mitochondrial membrane potential in the presence of substrates and after 1 mM GDP, determined from TMRM fluorescence shown in (f). (h) Oleate-induced depolarization expressed as Δψ decrease, determined from TMRM fluorescence. All values represent mean ± standard error (n = 5 for respiration, n = 3 for membrane potential). *p < 0.05, **p < 0.01, ***p < 0.001, **** p < 0.0001 versus Empty.

#### Bioenergetic effects of the presence of mouse-UCP1

Based on classical analyses of brown-fat mitochondria, the presence of functional UCP1 protein in mitochondria is expected to result in an elevated initial rate of oxygen consumption observed without further activation. It may be noted that this characteristic phenomenon can only be observed in isolated mitochondria, not in intact cells where this innate UCP1 activity is inhibited by cytosolic purine nucleotides.

We therefore performed respiration assays to determine the functional activity of the liver-expressed mouse-UCP1. In the presence of oxidizable substrates (glutamate, pyruvate and malate), mitochondria from control livers displayed a low initial rate of oxygen consumption (initial stretch of black line in Fig. 1c), whereas mitochondria from mouse-UCP1-expressing livers exhibited very significantly elevated basal oxygen consumption (initial stretch of green line in Fig. 1c) (Figure 1d, left panel). These experiments thus clearly demonstrated that innate uncoupling *is* a property of mUCP1 in mitochondria and is not caused by and dependent upon by factors specifically to be found in the brown adipose tissue environment, such as the mitochondrial membranes of that tissue.

#### Ectopically expressed mouse-UCP1 is sensitive to GDP

High innate thermogenic activity alone is not sufficient to confirm the proper functionality of the ectopically expressed mouse-UCP1. Inhibition of basal activity by GDP is generally considered a defining feature of functional UCP1 in brown-fat mitochondria (Cannon et al., 1973, Monemdjou et al., 1999, Shabalina et al., 2004, Oelkrug et al., 2010). Accordingly, we tested GDP sensitivity in mitochondria from AAV-mouse-UCP1 livers. Addition of 1 mM GDP did not have any effect in control liver mitochondria (black line in Fig. 1c) but in mouse-UCP1 mitochondria it very significantly inhibited basal respiration, down to exactly the level observed in control mitochondria (green line Fig. 1c; Fig. 1d). Thus, when expressed in the ectopic environment of the liver, mouse UCP1 retained both innate uncoupling activity and GDP sensitivity, and the fact that GDP could inhibit fully down to the basal level of respiration stresses the point that (mouse-)UCP1 is not “leaky”: the proton(-equivalent) conductance is totally abolished when mouse-UCP1 is GDP-inhibited (Shabalina et al., 2010). Correspondingly, the absence of innate proton conductance observed with UCP1 in reconstituted systems (liposomes and black lipid membrane and spheroplasts) must indicate that a (co)factor is missing in such experiments.

#### Ectopically expressed mouse UCP1 is oleate sensitive

Another defining feature of UCP1 is its ability to become (re)activated by fatty acids after it being inhibited by GDP inhibition (Rial et al., 1983, Shabalina et al., 2004, Gao et al., 2022, Shabalina et al., 2025). However, in contrast to the two distinct properties of innate activity and GDP inhibition, fatty acid activation is a relative property: the presence of UCP1 makes the mitochondria more sensitive to fatty acids (Shabalina et al., 2004), but all mitochondria can be uncoupled by fatty acids (Skulachev, 1991, Wojtczak and Schönfeld, 1993). In agreement with this, added oleate could uncouple control mitochondria (Fig. 1e). However, oleate much more potently induced oxygen consumption in GDP-inhibited mouse-UCP1 mitochondria (Figure 1c, e), confirming proper regulatory behavior of UCP1 when expressed in liver mitochondria.

To establish the Mitchellian behaviour of this liver UCP1 system, we also followed the effect of fatty acids on mitochondrial membrane potential with the fluorescence-based TMRM system where a quenched fluorescence indicate a higher membrane potential. As seen, compared with the control mitochondria, mouse-UCP1-expressing mitochondria showed reduced basal membrane potential, which was restored by GDP and subsequently decreased again upon oleate addition (Figure 1f). Quantification confirmed this regulatory pattern (Figures 1g and h).

When we plotted the membrane potential versus the oxygen consumption (Supplementary Fig. S1a), it became clear that the changes in oxygen consumption induced by oleate addition could be fully understood as being regulated in a Mitchellian way for the low oleate additions (left) but that the higher oleate amounts necessary to induce membrane depolarization also led to an inhibition of oxygen consumption capacity. Thus, the presence of mouse-UCP1 allowed for full uncoupling oleate effect within the tolerable oleate concentration window.

Taken together, these results demonstrate that ectopically expressed mouse-UCP1 in liver mitochondria exhibits the hallmark characteristics of native UCP1, including basal uncoupling, inhibition by GDP, and reactivation by fatty acids, thus confirming its functional integration and regulation. This opens for examining whether human-UCP1 behaves similarly to mouse-UCP1 – or species-specific characteristics exist.

### Expression of human UCP1 in liver mitochondria

#### Lack of impact of human-UCP1 on levels of mitochondrial oxphos proteins

To establish functional properties of human-UCP1, we injected mice with AAV-CRM8/TTR- human-UCP1, thus employing a liver-specific promoter, and analyzed UCP1 expression.

Relative human-UCP1 mRNA levels were well detected in AAV-injected mice (Figure 2a), at levels practically identical to those seen mouse-UCP1, principally facilitating functional comparisons between human-UCP1 and mouse-UCP1. Western blotting of isolated liver mitochondria revealed a band at ∼32 kDa corresponding to human UCP1, which matched the size observed in lysates from HEK293 cells transfected with human-UCP1 (Figure 2b, top row).

**Figure 2.**
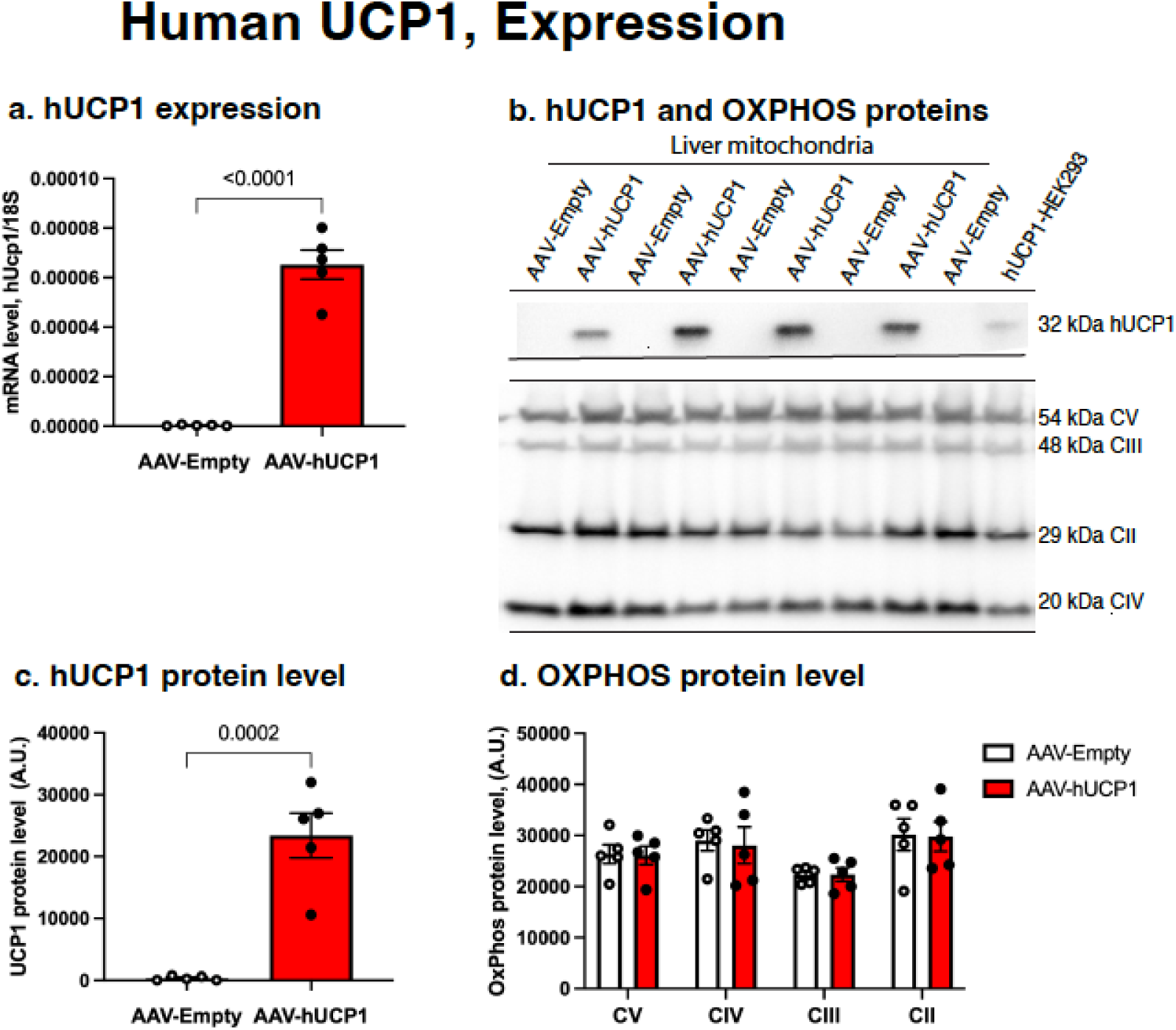
Expression of human-UCP1 and mitochondrial OXPHOS proteins. (a) Relative mRNA levels of hUCP1 in liver tissue of AAV-CRM8/TTR-hUCP1 or AAV-Empty-injected mice, normalized to 18S rRNA. (b) Western blot showing expression of hUCP1 in isolated liver mitochondria. Lysate from HEK293 cells expressing hUCP1 was used as a size reference (32 kDa) (upper panel). Representative western blot showing expression of mitochondrial oxidative phosphorylation (OXPHOS) complex subunits: ATP5A (Complex V), UQCRC2 (Complex III), SDHB (Complex II), and COX1 (Complex IV) (lower panel) (c, d) Quantification of UCP1 and OXPHOS proteins levels (n = 5 Empty and n = 4 hUCP1 liver mitochondria).

Previous studies have reported that ectopic overexpression of UCP1 can impair mitochondrial integrity, resulting in the depression of substrate oxidation, reduced cellular growth or in tissue autophagy (Bernal-Mizrachi et al., 2005, Keipert et al., 2009, Han et al., 2004, Xiong et al., 2021). Therefore, we evaluated whether UCP1 expression in the liver affected mitochondrial respiratory complexes. Western blotting for representative subunits of the oxidative phosphorylation system showed no difference in protein levels between mitochondria from AAV-hUCP1 and control (AAV-Empty) mice (Figure 2b, c). Thus, expression of human-UCP1 does not impair the abundance of major mitochondrial respiratory components.

#### Also human-UCP1 induces innate uncoupling in mouse liver mitochondria

In a previous study where human and rodent UCP1 were expressed in *Saccharomyces cerevisiae* it would seem that human-UCP1 lacked the high basal proton conductance observed in rodent UCP1; human-UCP1 showed activity only in the presence of fatty acids (Rodríguez-Sánchez and Rial, 2017). The authors concluded that while human-UCP1 is responsive to fatty acid activation and nucleotide inhibition, it lacks basal proton leak, suggesting a selective loss of intrinsic conductance in the human UCP1 ortholog. Another study, performed with mitochondria isolated from human brown adipose tissue depots, did, however, indicate that high innate activity could be observed, also with human-UCP1 (Porter et al., 2016).

We found that when oxygen consumption was measured in liver mitochondria with human-UCP1, the initial respiration in liver mitochondria was elevated compared to that in controls (Figure 3a, b). This was thus similar to the response observed with mouse-UCP1 mitochondria (Figure 1c, d). Thus, it can be concluded that human-UCP1 in this respect not is qualitatively different from mouse-UCP1, implying similar mechanisms for the activity (the proton(-equivalent) conductance).

**Figure 3.**
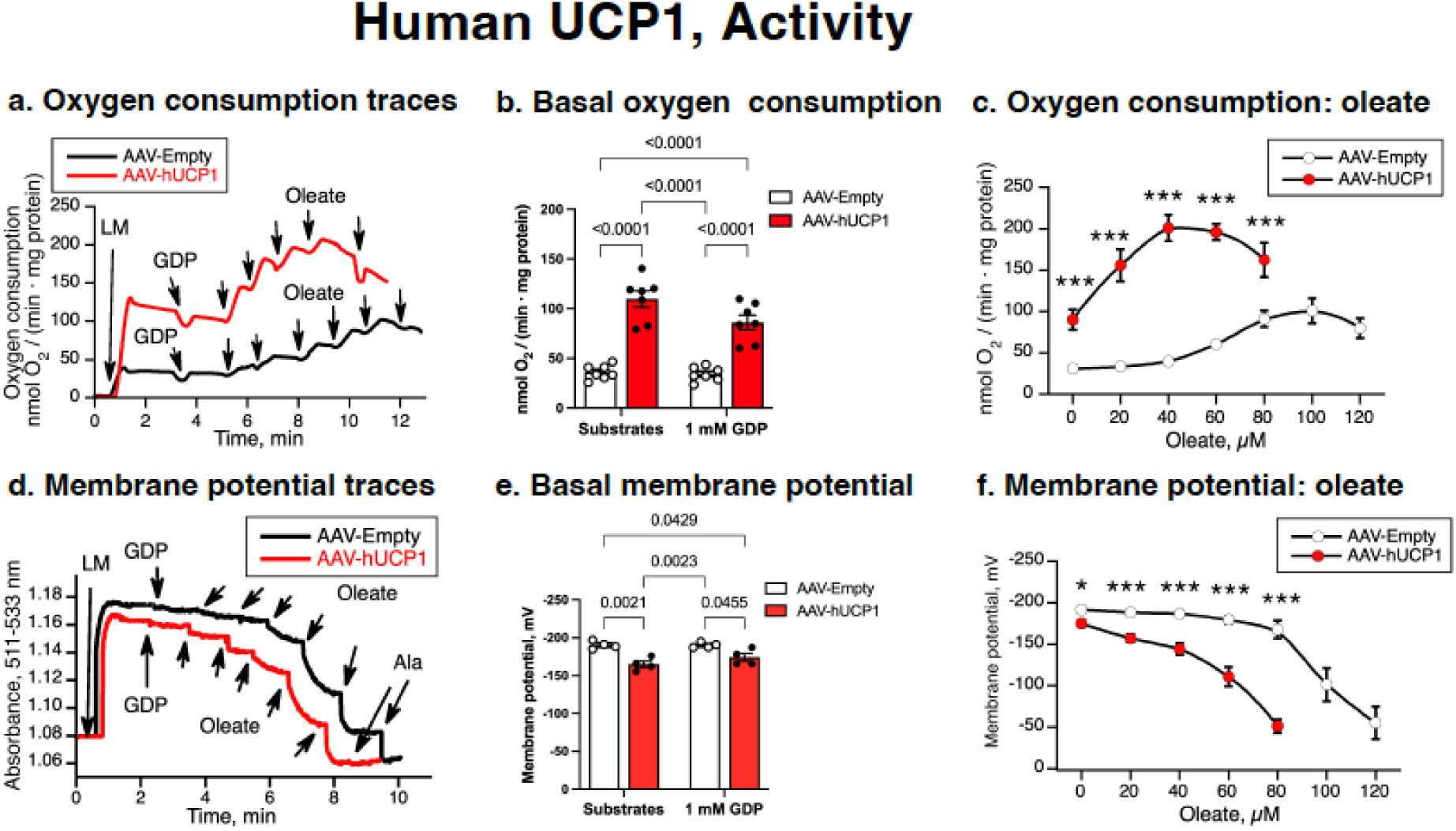
Human UCP1 activity in liver mitochondria. (a) Representative oxygen consumption traces in liver mitochondria (0.5 mg protein) expressing hUCP1 or control (shown in Figure 2), measured at 37 °C in the presence of malate, glutamate, and pyruvate, with 1 mM GDP and sequential addition of sodium oleate (20 µM). (b) Basal oxygen consumption rate (innate UCP1 activity) measured in the presence of substrates and after addition of 1 mM GDP compiled from (a). (c) Oleate-induced stimulation of respiration (GDP-inhibited rate subtracted) in hUCP1- expressing and control liver mitochondria. (d) Representative membrane potential traces recorded by safranin O absorbance at 511– 533 nm during oleate titration after 1 mM GDP. (e) Mitochondrial membrane potential in the presence of substrates and after GDP addition, quantified from the traces in (d). (f) Oleate-induced depolarization calculated from absorbance shift. All values are mean ± standard error (n = 7 for oxygen consumption, n = 4 for membrane potential). **p < 0.01, ***p < 0.001, **** p < 0.0001 versus Empty.

#### Ectopically expressed human-UCP1 is apparently GDP insensitive but is fatty acid-sensitive

However, concerning the second basic characteristic of mouse-UCP1 – the GDP-induced inhibition of activity, a key regulatory divergence emerged: addition of 1 mM GDP had only a minimal inhibitory effect on respiration in human-UCP1-expressing mitochondria, suggesting resistance to this canonical UCP1 inhibitor (Figure 3a, b). Initially this observation would seem to heavily question the functionality of human-UCP1 and/or make extrapolations from insights from mouse-UCP1 studies to human-UCP1 invalid.

In contrast, despite the human-UCP1 not being inhibited by GDP, addition of oleate effectively stimulated oxygen consumption (Figure 3c), consistent with the expected responsiveness to fatty acids. Again, the human-UCP1-containing mitochondria were much more sensitive to oleate than the control mitochondria (Figure 3c).

Membrane potential recordings enforced these findings. The human-UCP1-expressing mitochondria showed a reduced resting potential that was only slightly increased by GDP, but the mitochondria were strongly depolarized following oleate addition (Figure 3d). Quantification confirmed a persistently depolarized state with negligible GDP sensitivity and a robust response to fatty acids (Figure 3e, f). When oxygen consumption was plotted as a function of membrane potential (Supplementary Fig. S1b), a picture very similar to that obtained with mouse-UCP1 was obtained, with Mitchellian control of respiration at low levels of oleate addition but with respiration being inhibited by the oleate at higher oleate concentrations.

Taken together, these results demonstrate that human UCP1 expressed in liver mitochondria is functionally active and exhibits high sensitivity to fatty acids, but with nearly no sensitivity to GDP. This absence of the inhibitory GDP effect is disquieting, as it would seem to indicate that the human-UCP1 was overexpressed and inserted in such a way that control of proton conductance is lost, as has e.g. been observed with UCP3 expressed in yeast mitochondria (Harper et al., 2002). Such an uncontrolled proton leak could compromise tissue energy homeostasis, as previously observed in a vascular model of atherosclerosis (Bernal-Mizrachi et al., 2005). However, in our model, human-UCP1 did not reduce mitochondrial OXPHOS subunit levels (Figure 2), suggesting that there was no overt mitochondrial damage. We therefore proceeded to explore this regulatory divergence through detailed comparative analysis between mouse-UCP1 and human-UCP1.

### Human and mouse UCP1 exhibit species-specific regulatory sensitivities

For these direct comparative experiments, mouse-UCP1 and human-UCP1 were expressed under the same promoter (AAV8-CAG). To minimize possible influence from endogenous fatty acids we increased the BSA concentration in the isolation buffer from 0.2 % to 0.6 %. For these respiratory experiments, we employed a complete substrate mixture (pyruvate, glutamate, octanoyl-L-carnitine and malate).

To quantitate the apparent GDP insensitivity of human UCP1, we performed GDP titration experiments in mitochondria expressing either human-UCP1 or mouse-UCP1 (Figure 4a). As expected, mouse-UCP1-expressing mitochondria displayed potent GDP-induced inhibition of respiration; already the first addition of 0.3 mM GDP nearly fully inhibited the enhanced basal respiration induced by mouse-UCP1. In contrast, human-UCP1-expressing mitochondria exhibited minimal inhibition, even after cumulative GDP additions reaching 2.1 mM.

**Figure 4.**
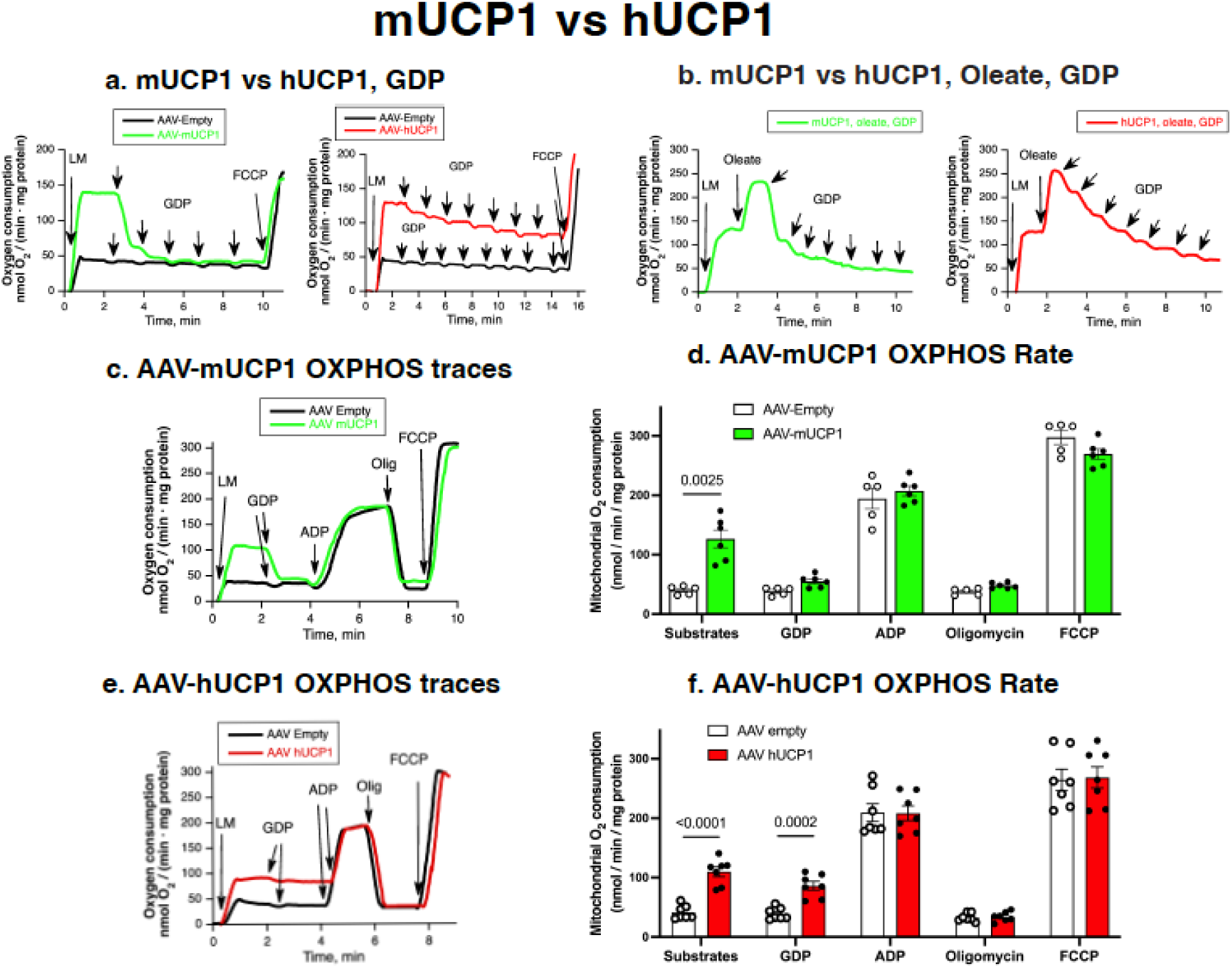
Differences in regulation of human versus mouse UCP1 in liver mitochondria. (a) Titration of oxygen consumption by GDP (each addition 0.3 mM) in liver mitochondria expressing mUCP1 or hUCP1 (both on AAV8-CAG promoter). The cumulative GDP concentration reached 1.5 mM for mUCP1 and 2.1 mM for hUCP1. At the end of the trace, 1.0 μM FCCP was added to assess oxidative capacity. (b) Titration of oxygen consumption by GDP (each addition 0.5 mM) in mUCP1 or hUCP1 liver mitochondria pre-stimulated by 50 µM oleate. The total addition of GDP was 3.5 mM in both groups. (c) Representative oxygen consumption traces in mUCP1-expressing mitochondria or control mitochondria under substrates (6 mM malate, 10 mM glutamate, 10 mM pyruvate, and 0.5 mM octanoyl-L-carnitine), 1 mM GDP, 1 mM ADP, 3 µg/mg oligomycin, and 0.8 µM FCCP. (d) GDP effect, OXPHOS capacity (state 3), State 4 (oligomycin-insensitive respiration), and maximal respiratory capacity (FCCP response) in mUCP1-expressing or control mitochondria. (e, f) Oxygen consumption traces and corresponding OXPHOS rates in hUCP1-expressing or control mitochondria. All values in (d) and (f) are mean ± standard error (n=5–7). P significance value is indicated on the graph.

When mitochondria were stimulated with 50 µM oleate, mouse-UCP1 mitochondria required higher GDP concentrations to achieve inhibition (Figure 4b), as expected (Shabalina et al., 2004). Concerning human-UCP1, it was clear that the enhanced respiration was not totally insensitive to GDP but human-UCP1 was evidently much less sensitive to GDP that was mouse-UCP1.

#### Oxidative phosphorylation is not impaired in liver mitochondria with human-UCP1

As it would seem that liver mitochondria carrying human-UCP1 were constantly somewhat uncoupled, the question was whether this condition would affect oxidative phosphorylation negatively and thus jeopardize the bioenergetic state of the liver cells. Oxidative phosphorylation was therefore assessed using sequential additions of substrates, GDP, ADP, oligomycin, and FCCP (Figures 4c and e).

In the mitochondria with mouse-UCP1, the initial high respiration was again fully inhibited by GDP. Addition of ADP induced, as expected, oxidative phosphorylation, and this process was inhibited by the ATP synthase inhibitor oligomycin. Remarkably, the presence of mouse-UCP1 did not qualitatively or quantitatively affect oxidative phosphorylation (Figure 4d).

In the mitochondria with human-UCP1, the initial high respiration was again totally insensitive to the GDP addition. After the addition of ADP, the respiration increased to exactly the same level as in control mitochondria, and even more remarkably, the effect of addition of oligomycin defined the entire enhanced respiration observed as originating from oxidative phosphorylation (Figure 4f). This must imply that the human-UCP1-induced uncoupled respiration was totally repressed as an effect of the initiation of oxidative phosphorylation.

A plausible explanation could be that ATP, produced during oxidative phosphorylation from exogenous ADP (and/or the added ADP itself), could act as inhibitor of human-UCP1.

#### ATP is a potent human-UCP1 inhibitor

To directly investigate the potency of ATP as a human-UCP1 inhibitor, mitochondria with human-UCP1 were subjected to titration with increasing concentrations of either GDP or ATP (Figure 5a) in respiratory studies. ATP did induce a concentration-dependent inhibition of human-UCP1 activity, with significantly greater potency than did GDP (Figure 5b). Thus, the apparent inability of GDP to inhibit human-UCP1 did not indicate that human-UCP1 displayed unregulated proton conductance; clearly human-UCP1 was responsive to purine nucleotide inhibition. The apparent IC₅₀ value for ATP (≈ 0.4 mM) was nearly 4-fold lower than that for GDP (≈ 1.4 mM) (Figure 5c).

**Figure 5.**
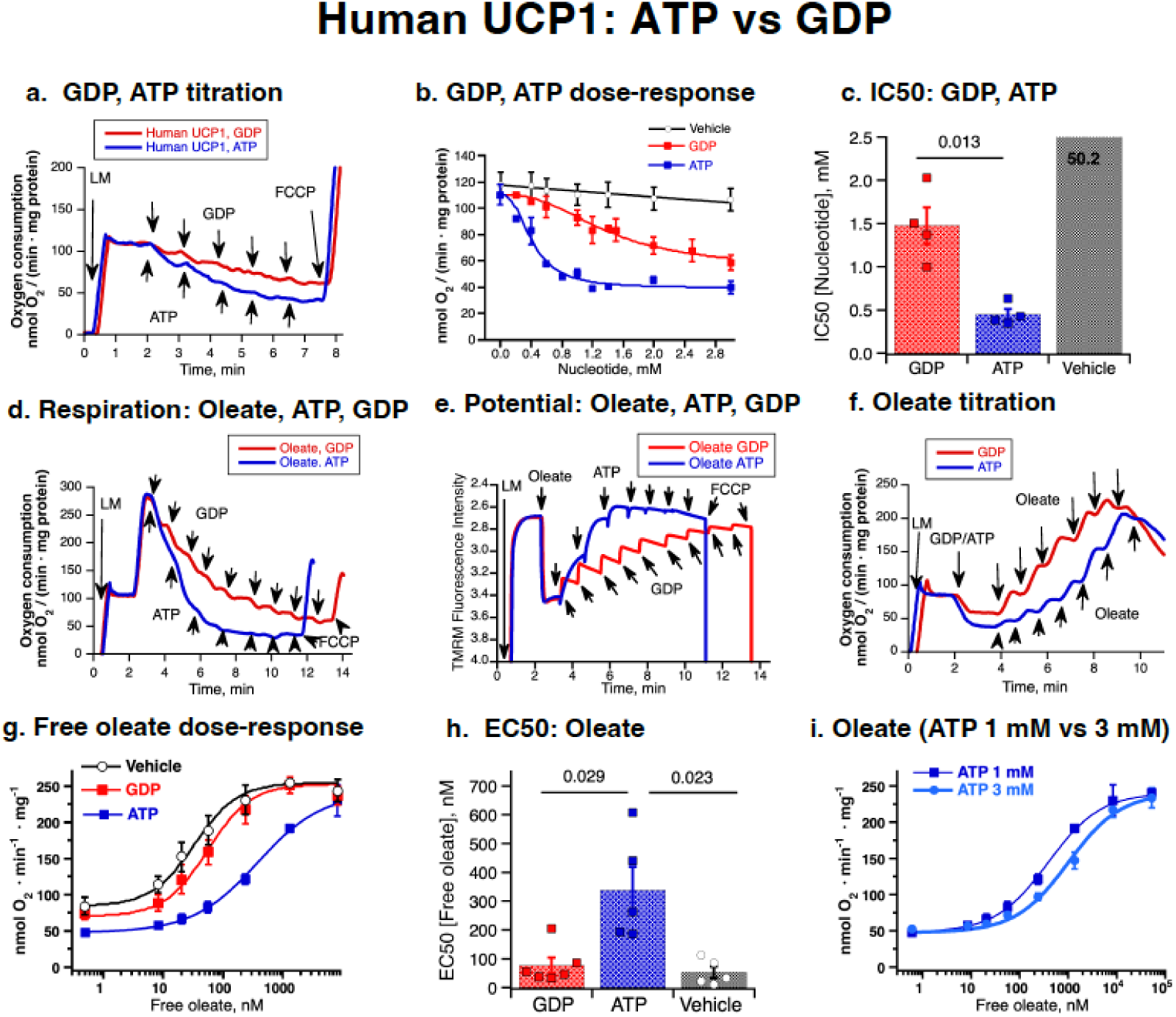
Differential inhibition of hUCP1 activity by ATP and GDP. (a) Representative oxygen consumption traces in mitochondria expressing hUCP1 during titration with increasing concentrations of GDP or ATP (each addition 0.5 mM). Full supply of substrates (6 mM malate, 10 mM glutamate, 10 mM pyruvate, and 0.5 mM octanoyl-L-carnitine), 3 µM CATR, and 3 µg/mg oligomycin were included. (b) Dose-response inhibition curves for GDP and ATP. (c) Half-maximal inhibitory concentrations (IC₅₀) for GDP and ATP determined by nonlinear regression. All values in b and c are mean ± standard error (n = 4 for each nucleotide and vehicle). P significance value is indicated on the graph. (d) Titration of oxygen consumption of hUCP1 mitochondria by GDP or ATP (each addition 0.5 mM) in hUCP1 mitochondria pre-stimulated by 50 µM oleate. Substrates and inhibitors were as in (a). (e) Titration of membrane potential (measured as TMRM fluorescence intensity in parallel to oxygen consumption shown in d) by GDP or ATP (each addition 0.5 mM) in hUCP1 mitochondria pre-stimulated by 50 µM oleate. All conditions and additions as in (d). (f) Oleate titration curve in hUCP1 mitochondria in the presence of 1 mM GDP or 1 mM ATP. Substrates and inhibitors were as in (a).(g) Oleate dose-response curves plotted against the calculated free oleate concentration. The corresponding dose-response curves plotted against the nominal (total added) oleate concentration are shown in Figure S2a. (h) EC₅₀ values for oleate-stimulated respiration in the presence of GDP, ATP, or the corresponding vehicle volume. All values in (g) and (h) are mean ± standard error (n=6 for GDP; n=5 for ATP and n=4 for vehicle). P significance value is indicated on the graph. (i) Oleate dose-response curves in the presence of 1 mM or 3 mM ATP. Experimental conditions were as in (f), except that ATP concentration was increased to 3 mM. All values are mean ± standard error (n=4 for ATP 1 mM and ATP 3 mM performed in parallel).

In mouse-UCP1, GDP not only inhibits innate proton conductance but also – in an apparently competitive way – the UCP1-dependent respiration induced by fatty acids (oleate) (Shabalina et al., 2004). The examine whether ATP in human-UCP1 also in these respects was equivalent to GDP in mouse-UCP1, we conducted a series of interaction studies.

We examined whether ATP (and GDP) could inhibit respiration in human-UCP1 mitochondria that had been pre-stimulated with 50 µM oleate. In this activated state, GDP again produced only modest inhibition, whereas ATP strongly suppressed respiration (Figure 5d). Membrane potential measurements performed in parallel confirmed that ATP effectively repolarized the mitochondria (Figure 5e).

To explore whether proton conductance inhibition through ATP could be overcome by fatty acid stimulation, similarly to what is seen in mouse brown-fat mitochondria (Shabalina et al., 2004), we performed titration by oleate in the presence of 1 mM GDP or 1 mM ATP (or without a nucleotide) (Figure 5f). We constructed dose–response curves for the nominal oleate concentration (Supplementary Fig. S2a) and for the free oleate concentration (Figure 5g). The presence of GDP had only a marginal effect on the ability of oleate to enhance respiration, but the presence of ATP substantially shifted oleate sensitivity (Figures 5f and Supplementary Fig. S2a). A direct comparison of EC₅₀ values revealed that significantly more oleate was required to overcome ATP inhibition than to overcome GDP inhibition (Figure 5h).

To determine whether higher nucleotide concentrations could lead to more potent inhibition, we increased the ATP and GDP titration to a final total of 3 mM (1 + 1 + 1 mM) (Figure S2b). The resulting respiration curves reflected the dose–response pattern observed for nucleotides in Figure 5b. We then followed this nucleotide titration with an oleate titration (Supplementary Fig. S2b). Notably, when mitochondria were pre-inhibited with 3 mM ATP, significantly more oleate was needed to reactivate respiration compared to the 1 mM ATP condition tested in parallel (Figure 5i), indicating a functional competition between ATP and fatty acids. In fact, the EC_50_ for oleate increased from 384 ± 21 nM free oleate in the presence of 1 mM ATP to 1000 ± 103 nM free oleate with 3 mM ATP.

These findings were verified by membrane potential recordings, which monitored mitochondrial polarization under the same conditions (Supplementary Fig. S2c).

These findings suggest that while fatty acid activation of human-UCP1 is robust, its nucleotide regulation is distinctly altered compared to mouse UCP1: ATP can potently inhibit human-UCP1-mediated uncoupling, whereas GDP is largely ineffective under the same conditions. Still, ATP possesses in this system all qualities observed for GDP in mouse-UCP1. The data therefore imply a functionally competitive interaction between nucleotides and fatty acids also for human-UCP1, similar to that observed in mouse-UCP1 (Shabalina et al., 2004).

### Molecular dynamics simulations pinpoint the structural alterations behind changed nucleotide sensitivity between mouse UCP1 and human UCP1

Our experimental results above demonstrate that GDP potently inhibits mouse UCP1 but shows poor inhibitory affinity toward human UCP1, whereas ATP does induce inhibition in human UCP1 (as it does in mouse UCP1, to be detailed elsewhere). To investigate the molecular basis underlying the difference in their responses towards nucleotides, we performed molecular dynamics simulations for the interaction of GDP with mouse UCP1 and human UCP1, as well as for the interaction of ATP with human-UCP1. For the starting configurations of ATP- human-UCP1 simulations, we chose to position ATP as in the ATP-bound human-UCP1 Cryo-EM structure (Kang and Chen, 2023). Since the conformation of GDP bound to UCP1 is unknown, we used three different starting configurations for each of the GDP-human-UCP1 and GDP-mouse-UCP1 simulations. In each case, the GDP molecule was initially placed slightly farther away than the ATP from the arginine triplet, with the phosphate group facing the triplet and the base oriented toward the intermembrane space. The starting configurations of all simulations are shown in Supplementary Fig. S3a.

To identify the primary nucleotide interaction sites, the fraction of total simulation time with interaction between ATP/GDP and human-UCP/mouse-UCP1 binding site residues was analyzed using ProLIF (Bouysset and Fiorucci, 2021), from which 16 residues were identified as constituting the nucleotide binding site (Figure 6a). Additonally, to analyze the interactions specifically between the nucleobases and UCP1, the distances between specific nucleobase atoms (indicated in Supplementary Fig. S3b) and the binding site residues were computed and presented as violin plots (Figure 6b).

**Figure 6.**
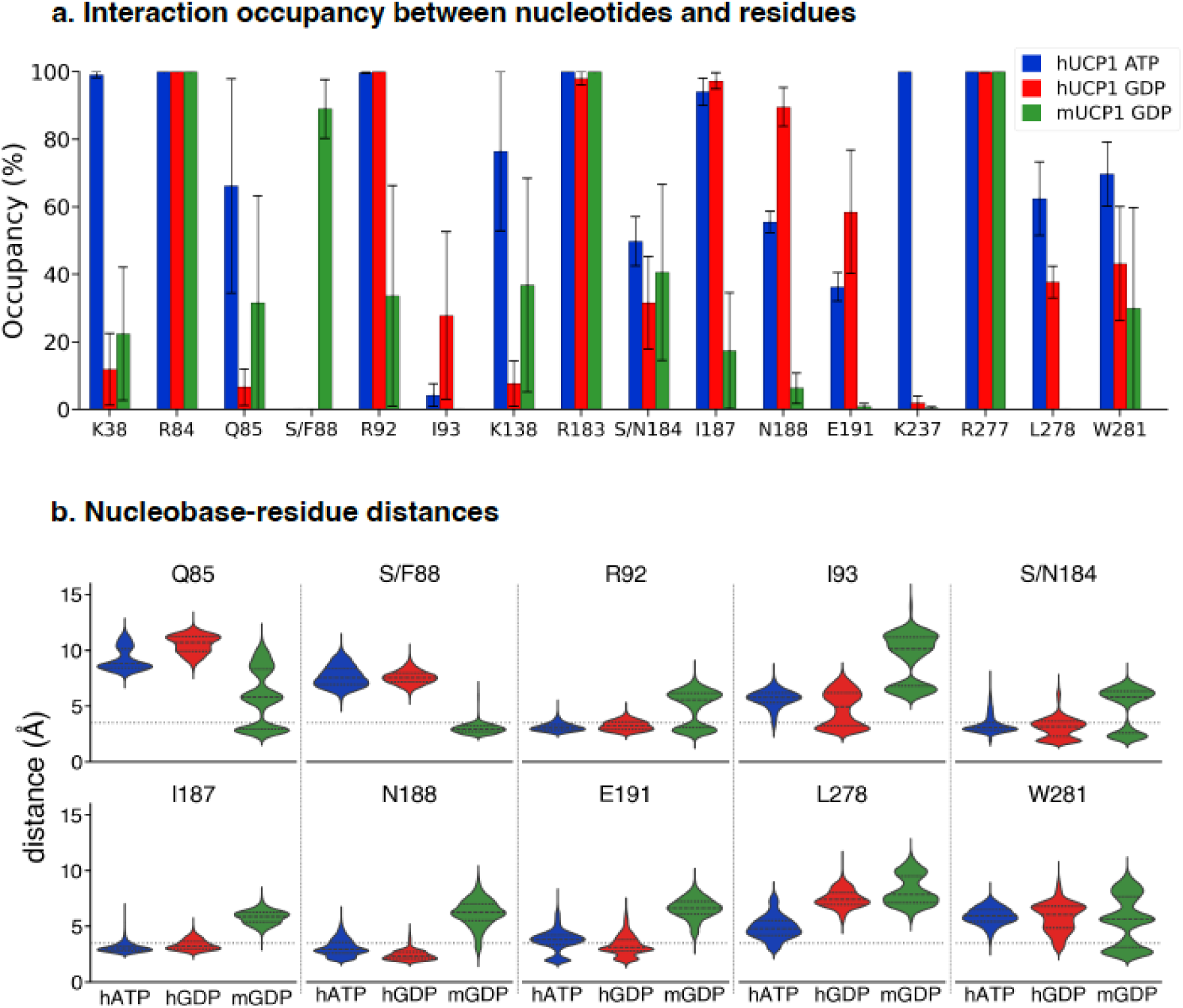
Nucleotide-residue interaction propensity and nucleobase-residue distances. (a) Fraction of simulation time in which interaction occurred between nucleotides and binding site residues. Only residues with an average fraction of time greater than 20 % in at least one simulation and a standard error smaller than the corresponding mean fraction are shown. Residues meeting these criteria are considered binding site residues. Only data from simulations that reached reproducible binding poses were included for GDP (hUCP1), n = 2; ATP (hUCP1), n = 3. The three different conformations were used for GDP (mUCP1), n = 3. Bars represent mean ± standard error. (b) Violin plots depicting the minimum distance between any of the green-colored nucleobase electronegative atoms (see Supplementary Fig. S3b) and UCP1 binding site residues (identified in a) during the last 300 ns of each trajectory (the arginine triplet residues and the matrix salt bridge residues are not shown as they interact with the phosphate groups, not the nucleobases). Black dashed lines within the violins indicate quartiles, and the grey dotted horizontal line at 3.5 Å indicates the defined threshold for identifying tight nucleobase- residue interaction.

#### The final ATP-bound conformation in ATP-human UCP1 simulations resembled that in the published cryo-EM structure

As the ATP was initially placed in the binding site identified by (Kang and Chen, 2023), it is unsurprising that in all three ATP-hUCP1 simulations, the final bound conformation closely resembled that in the corresponding Cryo-EM structure. The negatively charged phosphate groups were coordinated by the positively charged arginine triplet (R84, R183, R277), the matrix salt bridge lysine residues (K38, K138, and K237), and Q85 (Figures 6a and 7a). The adenine base formed cation-π interaction with R92, hydrophobic interactions with I187, as well as polar interactions with S184, N188 and E191 (Figures 6b and 7a). The sugar group formed hydrophobic contacts with L278 and W281. Additionally, R92 interacted with both the phosphate group and sugar moiety (Figure 7a).

**Figure 7.**
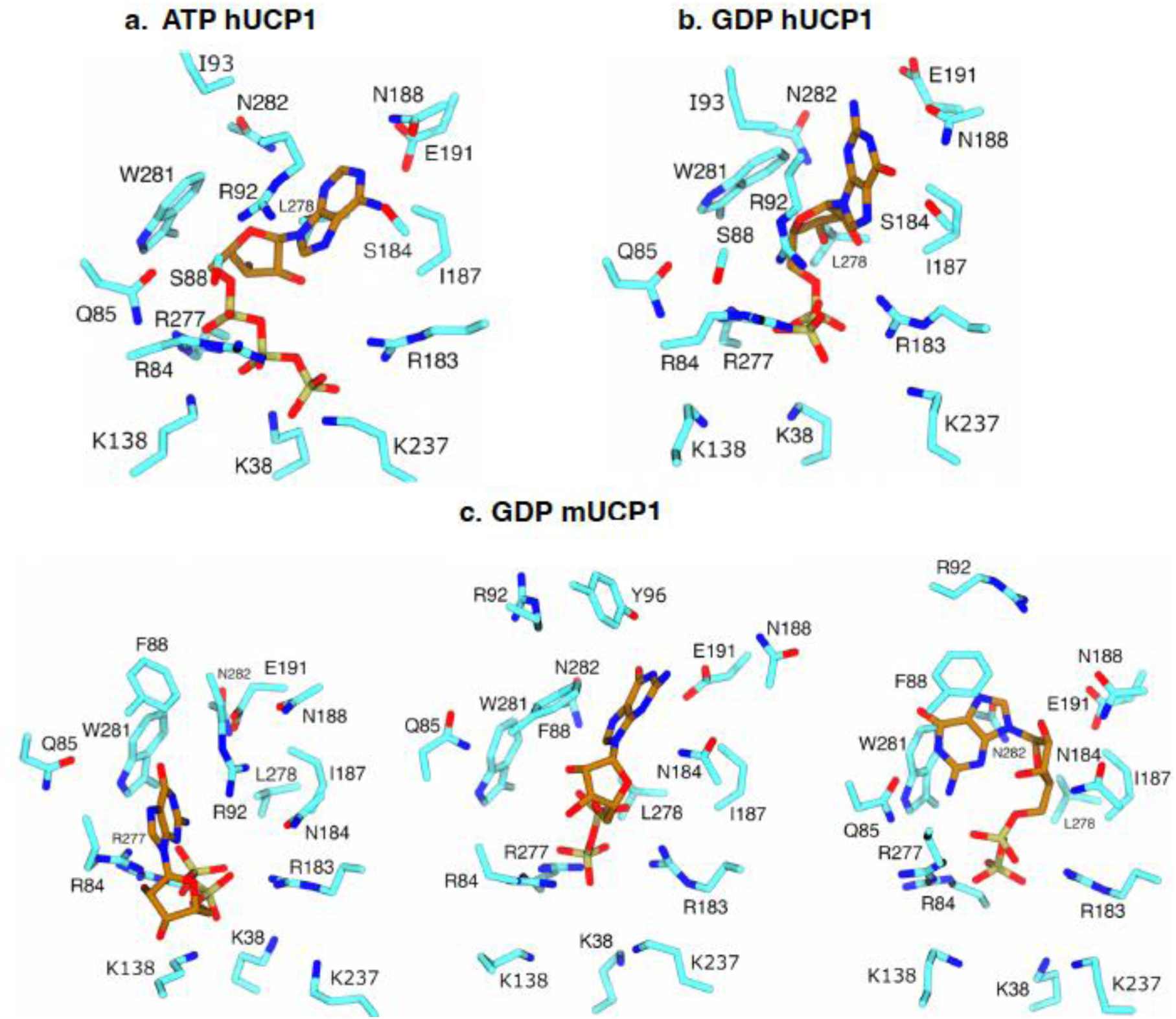
ATP/GDP-bound conformations of human-UCP1 and mouse-UCP1. (a) ATP-bound hUCP1. (b) GDP-bound hUCP1. (c) GDP-bound mUCP1. Residues and nucleotides shown as sticks and colored by their respective atom types (light blue: protein carbon, brown: nucleotide carbon, dark blue: nitrogen, red: oxygen). The bound conformations were determined using PyLipID (Song et al., 2022).

#### GDP binding to human-UCP1

Two of the GDP/human-UCP1 simulations converged to the same bound conformation, which at a first glance resembles how ATP bound human UCP1 (Figure 7b). However, GDP was much less likely to interact with all three matrix saltbridge network lysine residues (K38, K138, and K237), as well as to Q85 (Figure 6a and 7a). Only N188 showed a significantly elevated interaction occupancy to GDP relative to ATP in human-UCP1. This suggests that human UCP1 binds GDP more weakly than it binds ATP, despite an apparent similarity in their binding poses.

#### GDP binding to mouse-UCP1

The three mouse-UCP1/GDP simulations resulted in three distinct bound conformations. None of the three conformations resembled that observed for ATP in human-UCP1 (Figure 7a). However, all three GDP-mouse-UCP1 configurations feature stable nucleotide interactions with F88, which is absent in in either of the human-UCP1-nucleotide complexes (Figure 6a, b).

Inspection of the substrate-binding site of mouse-UCP1 and human-UCP1, alongside their corresponding sequences (Figure 8a), reveals that only two residues in the binding site differ: F88 and N184 in mouse-UCP1 are replaced by S88 and S184 in human-UCP1. The F88→S88 substitution introduces a substantial change, replacing a nonpolar bulky hydrophobic phenylalanine residue with a small polar hydrophilic serine. Indeed, our analysis shows that GDP and F88-mouse-UCP1 form persistent interactions, which are absent in human UCP1 (Figure 6a). In contrast, the N184→S184 substitution appears inconsequential for nucleotide binding. Therefore, it is likely that GDP’s reduced inhibitory affinity in human-UCP1 can largely be explained by the absence of F88, rather than by the absence of N184.

**Figure 8.**
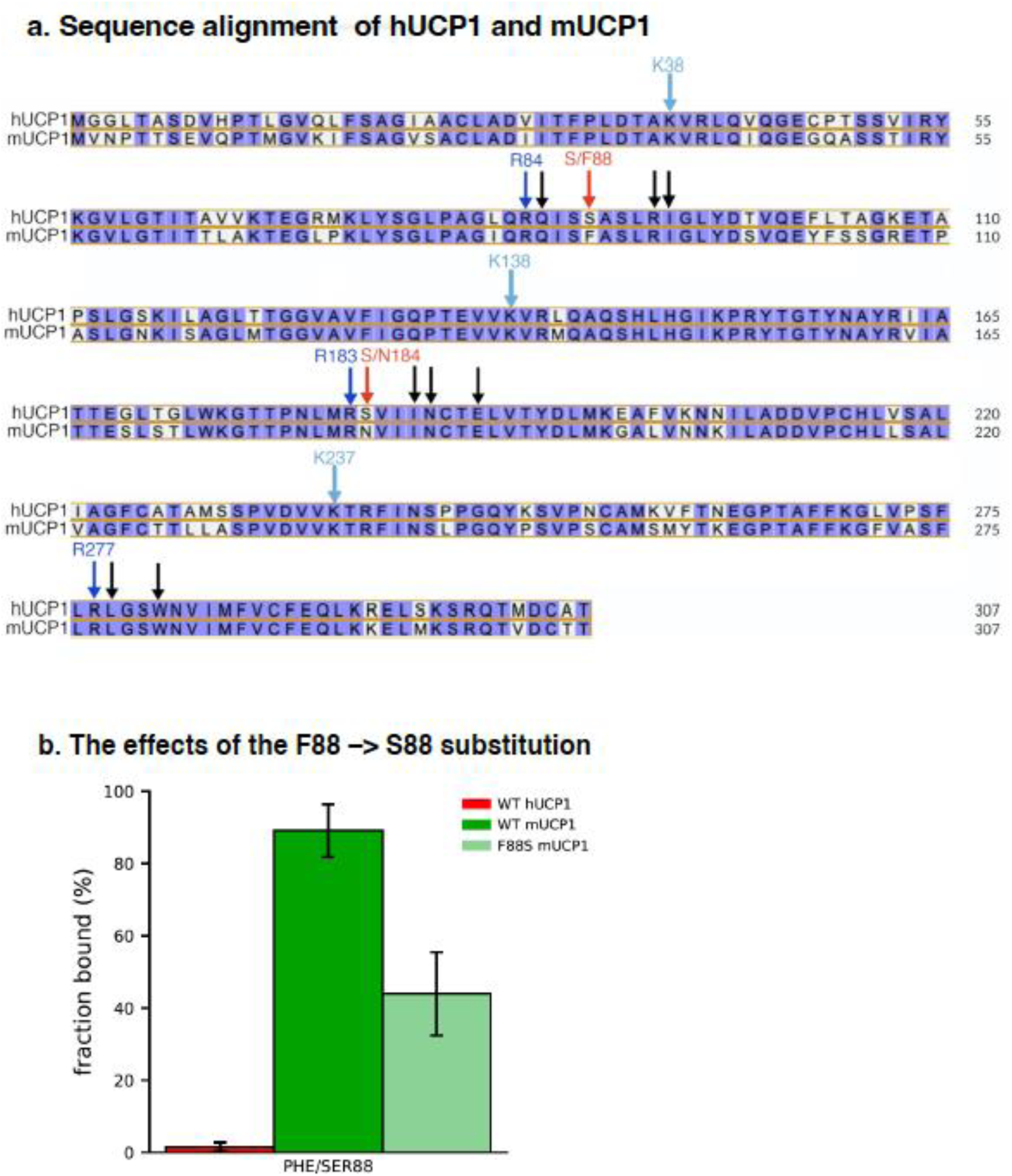
Protein sequence alignment and the effects of the F88 –> S88 substitution. a) Protein sequence alignment of hUCP1 and mUCP1. Arrows indicate residues in the nucleotide-binding site that interact with the nucleotides (Figure 6a): dark blue for arginine triplet residues, light blue for the matrix salt bridge lysine residues, and red for residues in the central binding site that differ between hUCP1 and mUCP1; black arrows for the rest of the residues in the binding site. b) Fraction of simulation time in which interaction occurred between GDP and residue 88 in wild-type human-UCP1, wild-type mouse-UCP1, and in-silico-mutated F88S mouse-UCP1. Introduction of F88S to mouse-UCP1 in the already bound state reduced interaction propensity between GDP and this residue during the following µsec.

#### F88 in mouse-UCP1 drives the high GDP-induced inhibition

To test the hypothesis that the F88→S88 substitution explains the loss of sensitivity to GDP observed in human versus mouse UCP1, we introduced the mutation F88S into mouse-UCP1 when the GDP was already in the bound position (after ≈ 1 µs) and simulated this mutated system for a further 1 µs (Figure 8b). Within this time frame, the mutation reduced the likelihood of GDP interaction with residue 88 by half (Figure 8b). Therefore, we suggest that the F88S substitution contributes to the reduced affinity of human UCP1 for GDP compared with mouse UCP1, thereby reducing its inhibitory potency.

## DISCUSSION

In the present study we have examined the function and regulation of human and mouse UCP1 in an ectopic system, i.e. UCP1 expressed in mouse liver and examined in liver mitochondria. We have found that the activity of UCP1 is well preserved in liver mitochondria, demonstrating that no additional brown-fat-specific factors are necessary for UCP1 activity. We have particularly found that not only mouse-UCP1 but also human-UCP1 display innate uncoupling – although this has earlier been doubted for human-UCP1. Both mouse-UCP1 and human- UCP1 also convey enhanced fatty acid sensitivity to the mitochondria, but they differ strongly in their sensitivity to purine nucleotides as inhibitors: GDP that is the classical UCP1 inhibitor for mouse-UCP1 is a very weak inhibitor for human-UCP1 – but ATP is well functional in human-UCP1. Fatty acid activation and ATP inhibition also interact similarly to what is established for GDP in mouse-UCP1.

Through molecular dynamics simulation we have arrived at a suggested structural explanation for the difference between human-UCP1 and mouse-UCP1. This difference seems to be largely explainable by a single amino acid mutation: from F88 in mouse-UCP1 to S88 in human-UCP1. The evolutionary role of this mutation can be discussed, and the analysis of human-UCP1 function in ectopic expression may have therapeutical implications.

### The study of ectopically expressed UCP1

The present investigation is evidently not the first to express UCP1 ectopically. In earlier studies, rodent UCP1 has been expressed e.g. in white adipose tissue (Kopecky et al., 1995, Yamada et al., 2006), in muscle and heart (Li et al., 2000, Couplan et al., 2002, Klaus et al., 2005) and in liver (Ishigaki et al., 2005). However, these earlier studies have often mainly aimed to establish systemic effects of the ectopically expressed UCP1, such as body weight loss, effect on muscle strength etc. When the resulting mitochondria have been studied, the bioenergetic characterization has been limited (e.g. rat-UCP1 (Couplan et al., 2002) and mouse-UCP1 (Keipert et al., 2009) in muscle mitochondria). We have here provided a detailed bioenergetic analysis of the effects of UCP1 on liver mitochondria. We have done this first with mouse- UCP1, so that the characteristics of UCP1 in the liver environment can be directly compared to those known from UCP1 in-situ studies. We conclude from those studies that the ectopically expressed mouse-UCP1 retains all the characteristics it displays in its native environment of brown-fat mitochondria: innate uncoupling, GDP inhibition, fatty acids stimulation and interaction between GDP and fatty acids. We have also done this with human-UCP1; this is the first characterization of human-UCP1 expressed in a mitochondrial system. We found that although there were species differences, the basic features of human-UCP1 did not deviate qualitatively from those of mouse-UCP1, except for a marked change in nucleotide sensitivity.

### Ectopically expressed UCP1 is innately thermogenically active

It was a clear outcome of our studies that even when ectopically expressed, the presence of UCP1 was associated with an enhanced basal respiration, the so-called innate uncoupling characteristics of UCP1. This is in clear contrast to the outcome when UCP1 has been incorporated into liposomes or black lipid membranes (Winkler and Klingenberg, 1994, Garlid et al., 1996, Urbánková et al., 2003, Cavalieri et al., 2022). (The absence of signs of this innate uncoupling in studies of cellular systems (not isolated mitochondria) such as yeast spheroplasts (Gagelin et al., 2023) or cell lines (Musiol et al., 2024), does not indicate that innate uncoupling is not a property of UCP1 but rather that in those systems UCP1 may be inhibited by cellular nucleotides.)

The implication of these observations is that a factor is found in mitochondria that is necessary for the manifestation of innate uncoupling. Endogenous fatty acids have been suggested as being the inducers of this activity. However, all earlier attempts to eliminate endogenous fatty acids by increasing BSA concentration, using cyclodextrins (lipid scavengers), or employing additional purification steps such as Percoll gradients, have consistently failed to suppress this UCP1-dependent uncoupling in isolated mitochondria (Rial et al., 2004, Shabalina et al., 2010, Yu et al., 2023, Shabalina et al., 2025). Also in the current studies of UCP1 in liver mitochondria we found that increasing the BSA concentration in the mitochondrial isolation buffer from 0.2% to 0.6% did not reduce mouse-UCP1 or human-UCP1 activity.

Another issue that has been discussed are the changes in brown-fat mitochondrial membrane lipids that are induced by acclimation to cold and have amply been described over the years (Cannon et al., 1975, Ricquier et al., 1975, Senault et al., 1975, Ocloo et al., 2007, Shimanaka et al., 2025). As the innate uncoupling is observable in liver mitochondria, the altered environments created by these phospholipid changes are clearly not essential for the activity of UCP1.

A general implication of these observations is thus that although UCP1 may be studied in e.g. black lipid membranes – with outcomes that are unnervingly close to those observed with the non-thermogenic UCP2 and UCP3 (Beck et al., 2007) – these observations may not be directly relevant for the understanding of how UCP1 functions in-situ.

### Also human-UCP1 displays innate uncoupling

We found that also human-UCP1 displays innate uncoupling. This clarifies an ongoing discussion in this respect. In a study where human and rodent UCP1 were expressed in yeast (Rodríguez-Sánchez and Rial, 2017), human-UCP1 showed activity only in the presence of fatty acids and lacked the high basal proton conductance observed with rodent UCP1. In contrast, in the only study to date that has addressed human-UCP1 in native human brown-fat mitochondria and directly compared its function to that in rodent brown-fat mitochondria (Porter et al., 2016), it was concluded that also human-UCP1 was associated with innate uncoupling. Our results adhere to that conclusion and imply that there is no qualitative difference between human-UCP1 and mouse-UCP1 in this respect.

### Human-UCP1 has practically no GDP sensitivity

The ability of GDP to inhibit UCP1 has until now been considered a basic property of UCP1, and this property was also what we observed here with mouse-UCP1 expressed in liver mitochondria, confirming that mouse-UCP1 was regulated similarly in its ectopic expression as in its native in-situ conformation. However, our experiments initially seemed to demonstrate what could have been a major qualitative difference between human-UCP1 and mouse-UCP1: an apparent lack of inhibitor sensitivity in human-UCP1. Still, it will be remembered that the use of GDP in connection with brown-fat bioenergetics is really a pragmatic choice. When it was realized that all purine nucleotide di- and triphosphates were able to inhibit UCP1 in isolated brown-fat mitochondria (Cannon et al., 1973, Nicholls et al., 1974, Huang and Klingenberg, 1995), it was considered most adequate experimentally to choose GDP rather than

ATP or ADP, as direct interaction with oxidative phosphorylation (and many other processes) in this way would be minimized. Correspondingly it was our examination of oxidative phosphorylation in human-UCP1-expressing mitochondria that made us realize that ATP had inhibitory power even for human-UCP1; a principally similar conclusion was reached by (Musiol et al., 2024), who observed that human-UCP1 in permeabilized HEK293 cells is less sensitive to GDP than to ADP. Thus, even human-UCP1 is nucleotide inhibitable – but not with all purine nucleotides, and this means that the general concept that “all” purine nucleotide di- and triphosphates are UCP1 inhibitors has to become reformulated: only adenine nucleotide di- and triphosphates are general UCP1 inhibitors. Still, the significance of the nucleotide as such – as compared to it being the phosphate groups that mainly mediate the binding – has been discussed (Jones et al., 2025).

### The binding site for the nucleotides

To identify which UCP1 residues were involved in nucleotide binding and through this to mechanistically understand why mouse-UCP1 but not human-UCP1 is GDP sensitive, we performed molecular dynamics simulations. Specifically, we examined how the interactions of human and mouse UCP1 with GDP differ, as well as how GDP binding is different from ATP binding. Recent structural and molecular dynamics studies have demonstrated nucleotide-dependent interactions and conformational dynamics within the central cavity of human UCP1 (Jones et al., 2023, Kang and Chen, 2023, Jacobsen et al., 2023, Gagelin et al., 2023, Jacobsen et al., 2025, Rathod et al., 2026). Regardless of species or the type of nucleotides, interactions between the nucleotide and the arginine triplet residues (R84, R183, R277) occur in virtually 100% of the time. In contrast, ATP and GDP interactions with UCP1 are distinguished by their propensity to bind the matrix salt bridge lysine residues (K38, K138, K237): while ATP almost always bind to all three residues, GDP rarely did.

ATP binding to human-UCP1 was stabilized by cation–π stacking with R92, polar interactions and hydrogen bonds to S184, N188, and E191, and hydrophobic interactions to I187, L278 and W281. ATP maintained a consistent position across replicas, matching that observed in the corresponding cryo-EM structure (PDB: 8hbw) (Kang and Chen, 2023).

In contrast, the simulations of human-UCP1-GDP complexes revealed that, while GDP can adopt a pose rather similar to that of ATP, charge interactions between its phosphate groups and matrix saltbridge network lysine residues K38, K138, K237 are drastically reduced compared to those of ATP, suggesting weaker binding.

In mouse-UCP1, however, residue F88 formed persistent interaction with deeply bound GDP, which might at least partly offset the loss of interaction with the matrix saltbridge lysine residues. Notably, in an MD simulation study of interaction between rat-UCP1 and GDP, residue F88 was also suggested as one of the residues important for nucleotide binding (Gagelin et al., 2023).

### A single mutation may be responsible for the loss of GDP sensitivity

Amongst those residues we have defined as binding residues there are only two that are different between mouse-UCP1 and human-UCP1: the F/S88 site and the N/S184 site (Fig. 8a). Concerning the N/S184 mutation, it is not functionally large (asparagine (N) is a small, polar residue, as is serine (S)), and the difference in occupancy or distance between the different formations is minor.

In contrast, the F88 -> S88 is structurally large (a nonpolar bulky hydrophobic phenylalanine residue as compared to a small polar hydrophilic serine) and it is associated with a qualitative difference in occupancy and in distance. Thus, we found it likely that this mutation was the culprit behind the loss of GDP sensitivity in human-UCP1. Indeed, mutating this phenylalanine residue in-silico to serine led to a rapid loss in occupancy – the GDP left the binding site.

### The evolutionary role of the F88S mutation

Analysis of UCP1 amino acid sequences clearly indicates that the F88 is the original one; it was present in the stem eutherian ancestor UCP1 (Keipert et al., 2024). The mutation to S88 would seem to have been introduced during primate evolution (and perhaps elsewhere as well). As any persistent mutation, it may be seen as a coincidence that was without marked phenotype effect – or as an evolutionary change that gave either potential problems or some fitness advantages to those that carried the mutation.

Initially it would sound very important that UCP1 through the F88S mutation has lost GDP sensitivity, - as exactly GDP sensitivity is a property we strongly associate with UCP1 function. An immediate presumption would be that this loss of GDP sensitivity would make human-UCP1 more easily activated in its native environment – as one of the factors that would inhibit it in competition against the stimulating fatty acids would then be less potent. However, it is normally considered that the levels of ATP and ADP are about one order of magnitude higher than those of GTP and GDP in the cell cytosol (Traut, 1994). Therefore, the loss of sensitivity to GDP would only very marginally affect the sensitivity of UCP1 to activating fatty acids. Still, one may consider primates an evolutionarily successful group, and the change may thus in some way have given the primates an advantage. However, given present information, the difference between human-UCP1 and mouse-UCP1 may just be an evolutionary coincidence, without any physiological significance.

### Possible therapeutical significance

The large bioenergetic effect of the F88S highlights the limitations of extrapolating rodent UCP1 data to human applications. This is especially important as there have been proposals to use ectopically expressed UCP1 therapeutically, in an attempt to diminish lipid accumulation systemically or in specific organs (such as liver to treat lipodystrophy (Ishigaki et al., 2005)). For immunological reasons it would be advantageous to use human-UCP1 for such treatments, but it is then important that the ectopically expressed UCP1 is under functional control. In this context, previous studies have reported that ectopic overexpression of UCP1 can impair mitochondrial integrity, resulting in reduced cellular growth or tissue damage (Han et al., 2004, Xiong et al., 2021, Bernal-Mizrachi et al., 2005). In UCP1-transgenic mice, regardless of the target tissue, some level of mitochondrial deficiency was observed. In transgenic skeletal muscle mitochondria expressing UCP1, a depression in substrate oxidation and reduced activity of mitochondrial complexes (Keipert et al., 2009) has been observed. High uncontrolled uncoupling, as observed in a vascular model of hUCP1 expression, resulted in increased blood pressure and accelerated atherosclerosis (Bernal-Mizrachi et al., 2005). However, our AAV-mediated expression appeared fully controlled. We did not observe mitochondrial impairment in liver mitochondria, neither in phosphorylation activity, oxidative capacity, nor in the levels of respiratory complexes, indicating that hUCP1 activity is conditionally regulated.

In conclusion, hUCP1 expressed in liver mitochondria functions as an active uncoupler with preserved fatty acid activation and ATP sensitivity but lacks classical GDP regulation. These findings provide crucial mechanistic insights and inform future strategies for the therapeutic exploitation of UCP1 in metabolic diseases.

## Supplementary Materials

Supplemental figures S1- S3 are provided.

## Author Contributions

IGS and JN conceived and designed the study. IGS performed mitochondrial experiments, analyzed data, and contributed to writing the manuscript. LJ conducted molecular simulations, performed analyses, and contributed together with ZWZ to manuscript writing. GRFB carried out AAV injections, phenotypic analysis of mice, mitochondrial experiments, and Western blotting. QN and JL performed Western blot analyses; JL and UA conducted mRNA quantification. BE designed the AAV constructs. HK supervised the molecular simulations. AE provided reagents and analytical tools. BC and AE contributed to data interpretation and analysis. JN also led the scientific discussion and critically revised the manuscript. All authors discussed the results and approved the final version of the manuscript.

## The Animal Ethics Board Statement

This study was approved by the Animal Ethics Board of the North Stockholm region and performed in accordance with national guidelines and regulations for the care and use of laboratory animals, in accordance with the guidelines of the European Communities Council Directive 2010/63/EU for the care and use of laboratory animals.

## Acknowledgments

We thank the Experimental Core Facility (ECF), Stockholm University, Sweden for technical assistance with the animals.

