## Supplementary Figures for "Differential Nucleotide Inhibition Profile of Mouse and Human UCP1 Expressed in Liver Mitochondria Is Associated with an F88S Mutation"

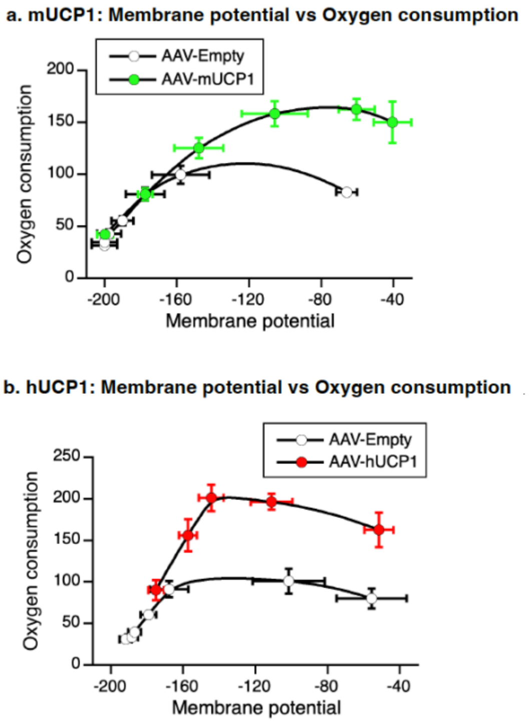


**Figure S1. Relationship between mitochondrial membrane potential and oxygen consumption in mUCP1- and hUCP1-expressing liver mitochondria.**
(a) Correlation between mitochondrial membrane potential and oxygen consumption during sequential titration with oleate in liver mitochondria expressing mouse UCP1 (mUCP1) or Empty control. Membrane potential values were derived from the experiments shown in Figure 1f–h, and oxygen consumption values from Figure 1c–e. Data are presented as mean ± standard error (n = 5 for respiration, n = 3 for membrane potential).
(b) Corresponding relationship for human UCP1 (hUCP1)-expressing liver mitochondria and Empty controls, calculated from the experiments shown in Figure 3. Data are presented as mean ± standard error (n = 7 for oxygen consumption, n = 4 for membrane potential).

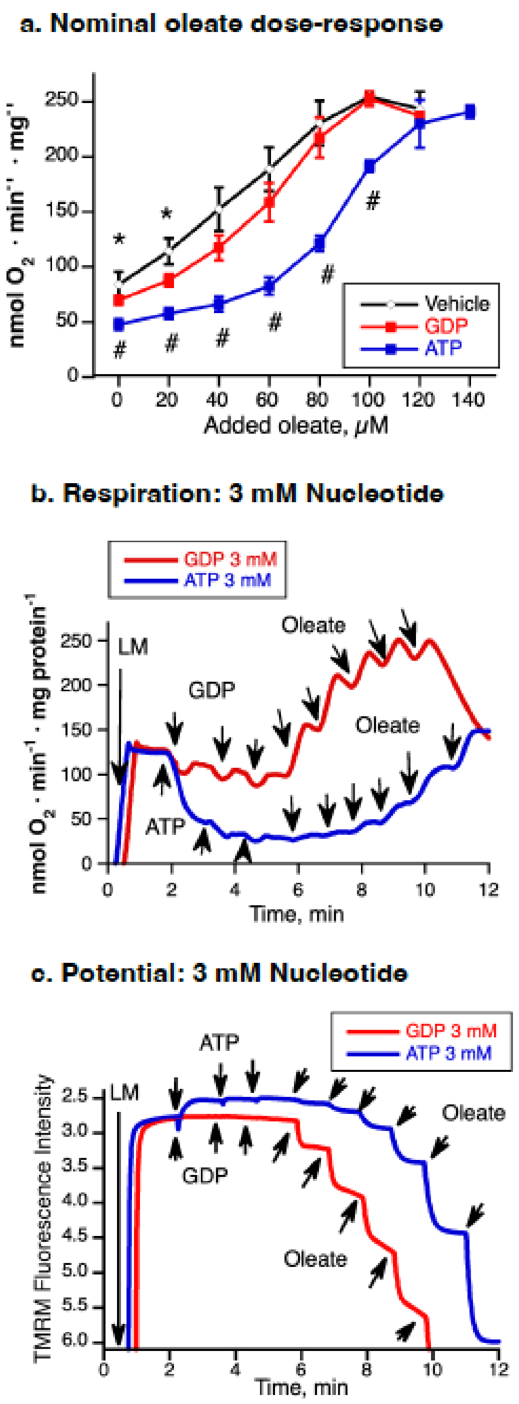


**Figure S2. Additional functional characterization of ATP- and GDP-mediated regulation of hUCP1.**

(a) Corresponding oleate dose-response curves from Figure 5g plotted against the nominal (total added) oleate concentration. * indicates a significant difference between GDP and vehicle; # indicates a significant difference between GDP and ATP.

(b) Representative oxygen consumption traces of hUCP1-expressing liver mitochondria during titration with GDP or ATP (1 mM each addition, 3 mM total), followed by sequential additions of oleate (20 µM each). All conditions and additions as Figure 5f except that GDP and ATP increased to 3 mM.

(c) Titration of membrane potential (measured as TMRM fluorescence intensity in parallel to oxygen consumption shown in a) by GDP or ATP (each addition 1 mM, 3 mM total) with the following titration by oleate (20 µM each addition). All conditions and additions as in S2b.

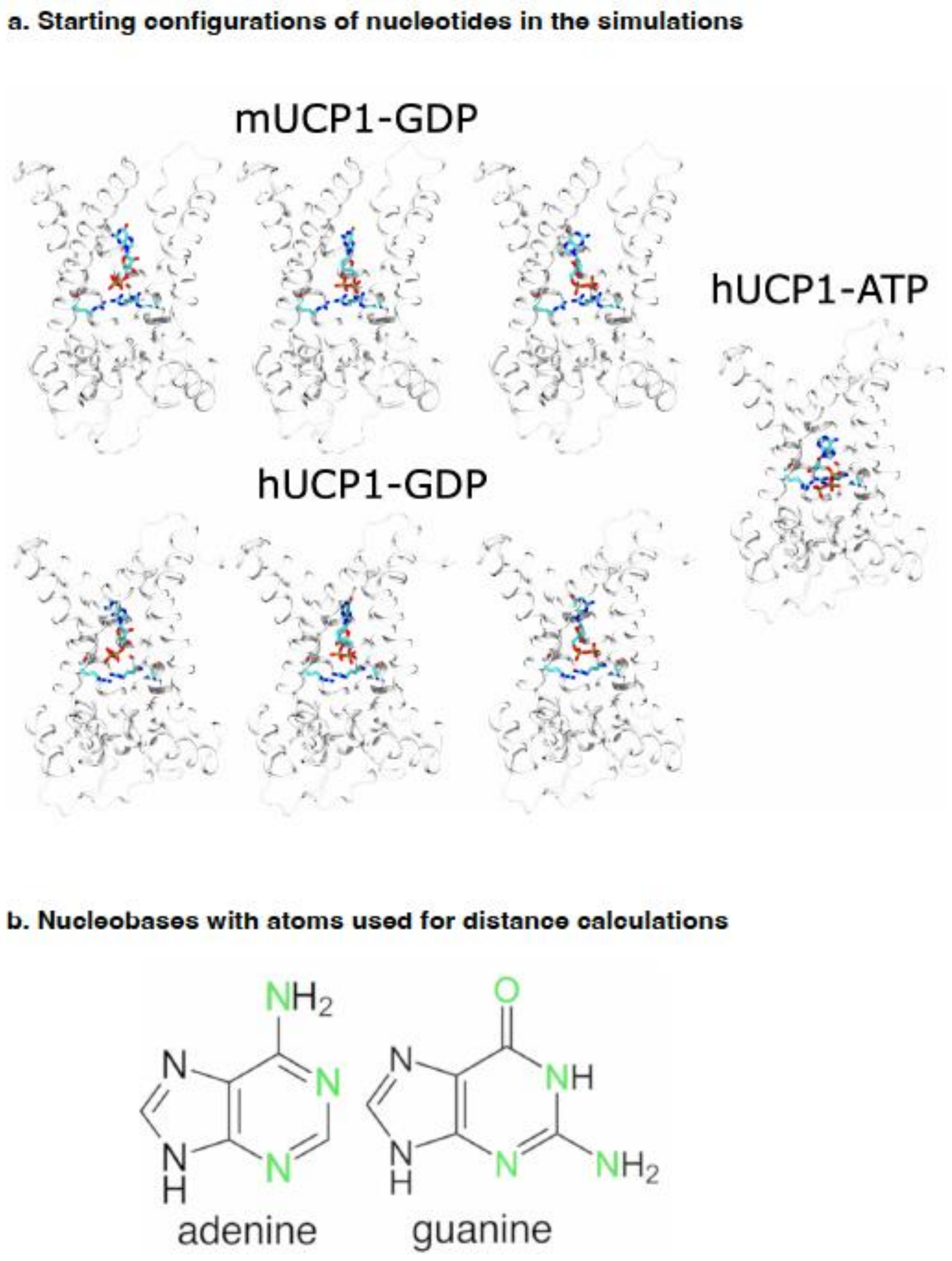


**Figure S3. Initial nucleotide configurations and nucleobase atom selection for molecular dynamics analyses.**
(a) Initial nucleotide configurations used for the molecular dynamics simulations. UCP1 is depicted as a white cartoon, with the arginine triplet (R84, R183, R277) and nucleotides shown as sticks and colored by atom type. Transmembrane helix 3 (TM3) is omitted for clarity.
(b) Nucleobases with the atoms used for the distance calculations in Figure 6b highlighted in green.
